# Signaling-dependent control of apical membrane size and self-renewal in rosette-stage human neuroepithelial stem cells

**DOI:** 10.1101/222125

**Authors:** Jan-Philip Medelnik, Kathleen Roensch, Satoshi Okawa, Antonio del Sol, Osvaldo Chara, Levan Mchedlishvili, Elly M. Tanaka

## Abstract

In the early developing nervous system, self-renewing neural stem cells are polarized and maintain an apical domain facing a central lumen. The presence of apical membrane is thought to have a profound influence on maintaining the stem cell state. With the onset of neurogenesis cells lose their polarization and the concomitant loss of the apical domain coincides with a loss of the stem cell identity. Very little is known about the molecular signals controlling apical membrane size. Here we use two neuroepithelial cell systems, one derived from regenerating axolotl spinal cord and the other from human ESCs to identify a conserved molecular signalling pathway initiated by lysophosphatidic acid (LPA) that controls apical membrane size and consequently controls and maintains epithelial organization and lumen size in neuroepithelial rosettes. This apical domain size increase occurs independently of effects on proliferation and involves a SRF-dependent transcriptional induction of junctional and apical membrane components.

## Introduction

Epithelial polarization and organization of a central lumen is a defining aspect of nervous system formation. During early vertebrate neurulation, neuroepithelial tissue, called the neural plate, is formed that subsequently undergoes folding to form the neural tube. At early stages, before and shortly after closure of the neural tube, the neuroepithelium consists of self-renewing, tight junction forming neural progenitor cells (NPCs) whose apical domains all face towards a central lumen/ventricle that contains the cerebrospinal fluid. At later stages, junctional complexes are remodeled; cells lose tight junctions and their polarized morphology, and at the same time onset of neuronal differentiation begins (Aaku-Saraste et al., 1996).

The capability of tight junction-containing neuroepithelium to organize into polarized, lumen-containing structures is also evident during neural organoid formation (Eiraku et al., 2011; Meinhardt et al., 2014) and in two-dimensional cultures of mouse E8 primary neural plate cell explants that spontaneously self-cluster to form two-dimensional mini-lumens termed “neural rosettes” (Elkabetz et al., 2008). Neural rosettes consist of small cellular units in which the neuroepithelial cells all orient their apical domain to a common lumen-like center. Due to their self-organizing capacity neural rosettes can be regarded as in-vitro, two-dimensional versions of the neural tube. Similar to the gradual loss of neural stem cell identity in the developing neural tube, the ability of neuroepithelial cell explants derived from E8 mouse embryos to organize in rosettes is also restricted to a short window of early neural development prior to down-regulation of tight junctions and a loss of polarization. It has been difficult to maintain rosettes in culture as they tend to spontaneously progress toward more differentiated, less polarized phenotypes (Elkabetz et al., 2008).

An alternative source of rosette-forming-, tight junction-containing neuroepithelium has been mouse and human ES cell cultures early after neural lineage induction. The early rosette-forming neuroepithelial state is considered an important state for stem cell differentiation, as it represents the early neural plate- or neural tube stage at which cells have committed to the neural lineage but are still susceptible to instructive cues that pattern the CNS along the anterior/posterior and dorsal/ventral axes (Broccoli et al., 2014; Elkabetz et al., 2008; Li et al., 2005; Pankratz et al., 2007). In human ES cells, dual inhibition of SMAD signaling by noggin/dorsomorphin and SB431542 efficiently directs cells along the neural lineage and results in rosette formation (Chambers et al., 2009; Zhou et al., 2010). However, upon passaging, cells successively form smaller rosettes in which cell polarization and tight junctions get reduced over time, until the neuroepithelial cell state is completely lost. Elkabetz *et al*. reported that addition of SHH and the Notch signaling agonists Delta and Jagged-1 during plating of rosette forming NPCs could prolong the rosette forming state for approximately one passage by presumably promoting symmetric-self-renewing cell divisions (Elkabetz et al., 2008). Few studies have identified signaling factors that control growth and expansion of the luminal neural tube area independent of cell proliferation and beyond the factors mentioned above, there are no known factors that specifically sustain a large apical domain- and rosette forming neuroepithelial state. Identification of factors that can sustain/promote neuroepithelial lumen formation independently of or in parallel to symmetric, self-renewing divisions should have a profound effect on propagating the neural rosette forming state, and might forestall the timely differentiation to later stage neural progenitors/committed cells. Interestingly, the sustained self-renewal of early neuroepithelial cells is a characteristic feature of spinal cord regeneration in the axolotl in which spinal cord lesion induces cells to enter the neuroepithelial state. This state is sustained and self-renews for at least two weeks to yield an outgrowth of the severed spinal cord tube (Rodrigo Albors et al., 2015; Rost et al., 2016).

Given the importance of the tight junction-containing early neuroepithelial state during neural tube formation and neural tube regeneration in the axolotl we sought to identify a molecular factor that promotes expansion of the apical lumen in neuroepithelial rosette cells. Here we have identified lysophosphatidic acid (LPA) as a molecule which can cause an expansion of the apical domain of axolotl and human neuroepithelial cells, and growth of rosette luminal area independent of cell proliferation. Continuous treatment of HESC-derived neuroepithelial cells with LPA yields the propagation of the rosette forming state for at least 52 passages. Correspondingly, we see a suppression of neuronal differentiation and sustenance of the SOX2^+^, proliferative state. LPA acts through Rho and SRF to promote upregulation of tight junction- actin-related- and apically-associated proteins.

## Results

### Addition of serum or LPA to axolotl and human ESC-derived neuroepithelia induces rosette enlargement

Axolotl spinal cord regeneration is a unique system in which an injury-initiated wound response induces the self-renewal and expansion of neuroepithelial cells that grow a lumen containing neural tube which can recapitulate neural patterning (Rodrigo Albors et al., 2015; Tanaka and Ferretti, 2009; Walder et al., 2003). The lumen of the neural tube is a closed structure that is entirely surrounded by the apical cell surfaces of the neuroepithelial cells that outline the lumen. A key extracellular mediator of the injury response is serum, which represents the liquid component of clotted blood. We therefore asked if serum and any of its major components could induce an expansion of neuroepithelial rosette formation in cultured axolotl spinal cord cells. While the two-dimensionality and their open structure neural rosettes do not represent closed lumina, they still represent lumen- or neural tube-like structures that recapitulate many features of a true lumen and thus can be used as a cell culture-based system to identify factors effecting lumen organization. We exposed axolotl spinal cord-derived SOX2^+^ neuroepithelial cell explants to a three day pulse of serum corresponding to the injury kinetics *in vivo* (Rodrigo Albors et al., 2015; Sugiura et al., 2016). Strikingly we observed the organization of the cells into larger rosette structures over time (Figure 1A, upper left and middle panel). To identify the serum component responsible for this effect, we assayed LPA which is a signaling phospholipid that is a major signaling molecule in serum. When we exposed cells to concentrations of LPA in a similar range compared to those measured in serum (Yung et al., 2014) we recapitulated the larger rosette phenotype similar to the phenotype observed with serum treatment (Figure 1A, upper right panel). We concluded from these observations that LPA is a serum-borne signaling molecule that can induce neuroepithelial rosette enlargement in axolotl spinal cord NPCs. We next asked if the action of LPA is unique to axolotl neuroepithelia, or whether it represents a conserved signaling pathway that can regulate rosette lumen size in other vertebrate neuroepithelia. Furthermore, we sought an established cell culture system as an alternative to the axolotl primary cell culture system to further study the action of LPA on lumen organization. We therefore asked if serum- and LPA induced an increase in rosette size in human ESC-derived neuroepithelial cells. We implemented a modified version of a three-dimensional Dual SMAD signaling inhibition protocol to efficiently differentiate HES cells into rosette-forming NPCs (Khattak et al., 2015). In these cultures more than 85% of NPCs expressed the neuroectodermal transcription factor SOX1. These cells organized in small rosettes and expressed the tight junction-associated protein ZO-1 at their luminal side. SOX1^+^ NPCs could be maintained for at least 10 passages without losing SOX1 expression and their small rosette organization when cultured in N2B27 + FGF2/EGF medium (Figure 1A, lower left panel, Figure S1). Upon incubation with 20% serum or 15 μM LPA, the human NPCs reorganized to form larger rosettes within 18 – 24 h indicating that the signaling pathway leading to the formation of larger rosette structures is conserved between axolotl- and human NSCs (Figure 1A, lower middle and right panel).

**Figure 1.**
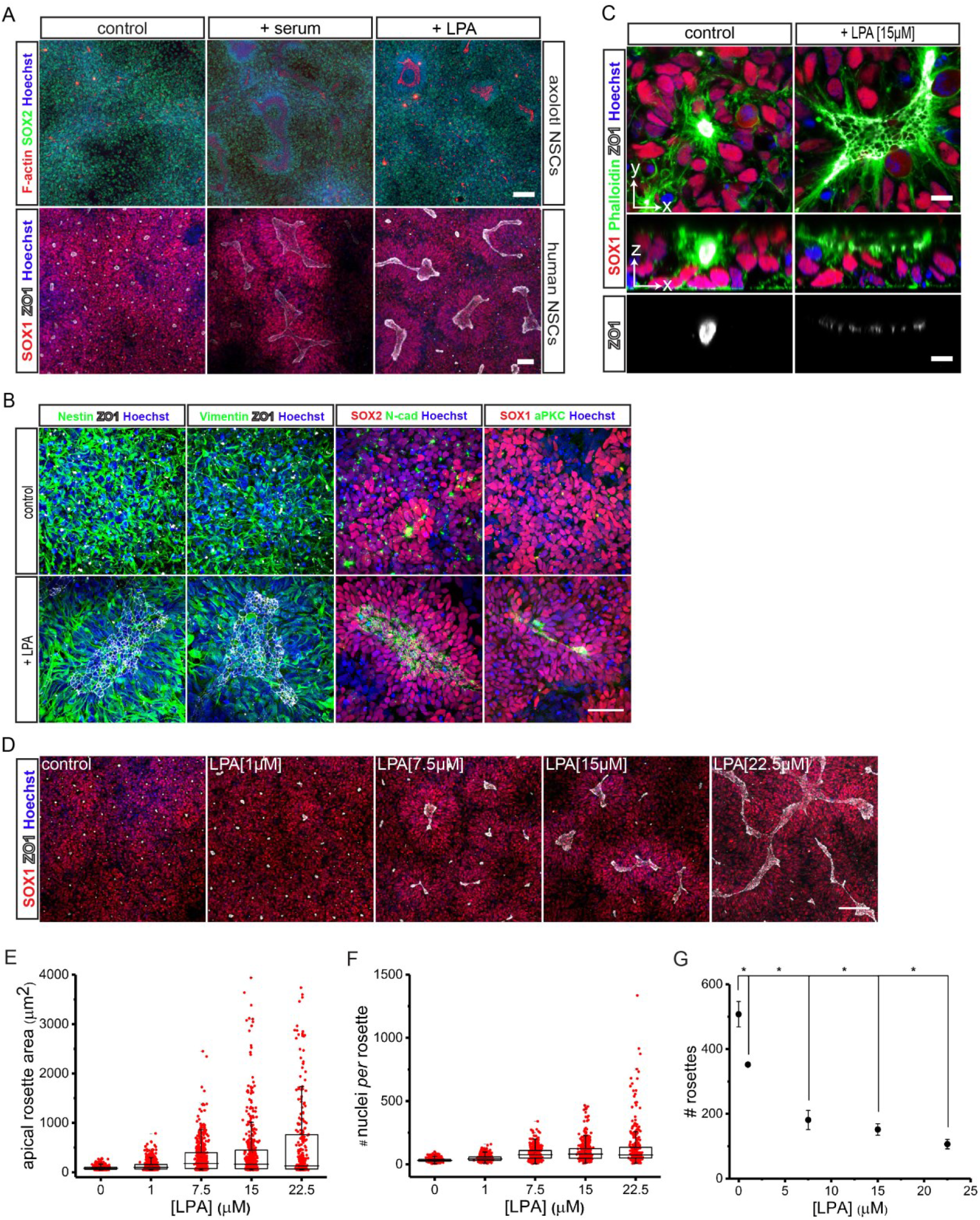
Serum and LPA induce large rosette formation in cultured axolotl- and hESC-derived NPCs. (A) Immunostaining of axolotl- and hESC-derived NPCs for SOX2 (upper panels), SOX1 and ZO-1 (lower panels) shows induction of large rosette formation in the presence of serum or LPA. F-actin in axolotl NPCs was stained with Alexa Fluor 647 Phalloidin. Scale bars are 200 μm (upper panels) and 100 μm (lower panels). (B and C) Immunostaining of HESC-derived NPCs for Nestin, Vimentin, SOX1, SOX2, N-cadherin, aPKC and ZO-1. F-actin was labeled with Alexa Fluor Phalloidin 488. Y-x-images are top view- and z-x-images side view images. Scale bars are 50 μm (B) and 10 μm (C). (D) Immunostaining of NPCs that have formed enlarged rosettes after a 24 h exposure to different LPA concentrations (0 – 22.5 μM) for SOX1 and ZO-1. Scale bar 100 μm. (E,F,G) Quantification of the frequencies of the apical rosette lumen size areas in μm^2^ at different LPA levels. Red dots indicate individual apical rosette sizes (E), # of SOX1^+^ cells participating in rosette formation (F) and mean # of rosettes per analyzed image G). Statistical analysis was performed using an ANOVA test followed by a Tukey a posteriori test. Data represent mean of medians ± SEM, *= *p* < 0.05. *n* = 3. Nuclei were labeled with Hoechst 33342. See also Figures S1 and S2.

### HESC-derived rosettes express early neuroepithelial markers and exhibit apical to basal polarity

We next characterized the HESC-derived neural rosette cultures and their change in organization in response to LPA. Rosette-NPCs are considered to represent a neural stem cell type whose lumen-organizing capacity and neuroepithelial marker expression strongly resembles early neuroepithelium at the neural plate stage (Elkabetz et al., 2008; Li et al., 2005). We compared the neuroepithelial- and junctional marker expression between control and LPA-treated, large rosette NPCs. Both control-small and LPA-induced large neural rosette NPCs expressed the junctional marker ZO-1, the intermediate filament proteins Nestin and Vimentin and the neuroectodermal transcription factors SOX1 and SOX2 (Figure 1B). Consistent with their ability to organize a lumen-like structure, the rosette-NPCs in both conditions expressed the apically localized proteins N-cadherin, aPKC and CD133 and showed enriched localization of F-actin at their apical side, indicating the strong apicolateral polarization of the cells toward the rosette center (Figures 1B and 1C, Figure S2). While the control- and LPA-treated rosettes expressed a similar spectrum of neuroepithelial- and polarity markers we observed striking differences in the spatial organization of the cells. In control rosettes the lumen exhibited a strongly constricted morphology with clustering of ZO-1 immunostaining so that individual cell-cell junctions were not resolvable at the magnification that we normally employed (Figure 1C, upper left panel). XZ scans of a rosette structure showed the tight clustering of ZO-1 into a defined point, reflecting the small area of apical domain in the control NPCs (Figure 1C, lower and middle left panel). In contrast, in the presence of LPA, the luminal surface had become much larger and cells adopted an unconstricted morphology in which the apical domains of the cells had become broader and where the individual cell-cell junctions were easily recognizable by staining for ZO-1 (Figure 1C, right panels).

### LPA increases rosette lumen size in a concentration-dependent manner

We next tested if the LPA-induced rosette size increase could be regulated by incubating the cells with different LPA concentrations. Exposure of NPCs to different LPA doses over a period of 24 h and immunostaining of the cells for SOX1 and ZO-1 to visualize the extent of rosette formation resulted in the formation of larger rosettes with larger luminal surfaces in a concentration-dependent manner (Figure 1D). We defined the lumen surface area of a rosette as the entire ZO-1-positive area completely enclosed by SOX1^+^ nuclei. Quantification of the LPA-dependent rosette lumen area revealed that distributions of apical rosette-lumen area are shifted towards larger values. In particular, lower quartiles, upper quartiles and interquartile ranges monotonically increase from 59.4, 101.6, 42.2 μm^2^ in control NPCs to 64.1, 760.9, 696.9 μm^2^, respectively, at 22.5 μM LPA (Figure 1E). We next quantified the number of SOX1^+^ cells per rosette. Analogously with rosette-lumen area, lower quartiles, upper quartiles and interquartile ranges monotonically increase from 24, 40 and 16 in control NPCs to 50, 134, 84, respectively, at 22.5 μM LPA (Figure 1F). Concomitant to the increase in rosette size the total number of rosettes quantified per image decreased from a mean of 508 ± 39 rosettes in the control to a mean of 106 ± 15 rosettes at a concentration of 22.5 μM LPA (Figure 1G). Altogether the data indicate that LPA increases rosette size in a concentration-dependent manner and that large rosette formation results from an increase in the total rosette lumen surface area and an increase in the number of cells per rosette.

### Large rosettes can be induced in the absence of cell proliferation

Our results above showed that large rosette formation coincides with an increase in the mean number of SOX1^+^ cells that surround a lumen. We wanted to determine whether the average increase in the cell number per rosette in the presence of LPA could solely be attributed to an increase in cell proliferation. Human ES cell- and iPSC-derived NPCs have been reported to either increase- or decrease their proliferative activity in response to LPA. The opposing results were attributed to differences in the cell origin and the culturing condition (Frisca et al., 2013; Hurst et al., 2008; Pebay et al., 2007). As NPCs progress along lineage differentiation their responsiveness to LPA likely changes. The action of LPA on defined rosette-forming neuroepithelial cultures as opposed to other types of NPC cultures has not yet been adequately clarified. We therefore measured proliferative activity in our cultures by exposing the NPCs to LPA concentrations ranging from 0.5 μM to 2.5 μM and adding the thymidine analogue 5-ethynyl-2’-deoxyuridine (EdU) which labels cells undergoing DNA synthesis 45 min prior to fixation. Quantification of the percentage of EdU^+^/SOX1^+^-cells revealed a 66 % increase (25 ± 1 % EdU^+^/SOX1^+^-control NPCs vs. 41.5 ± 0.4 % EdU^+^/SOX1^+^-LPA-treated NPCs) in the proliferative activity of NPCs that were exposed to 0.5 μM LPA compared to the control (Figures 2A and 2B). Concentrations higher than 0.5 μM did not further increase the proliferation in our experiment (data not shown) which already suggested that the dose-dependent increase in rosette size is unlikely to be related to proliferation. To further investigate the importance of cell proliferation on large rosette formation we sought to block the proliferative activity of the NPCs during the LPA treatment with hydroxyurea (HU) (Koc et al., 2004). To ensure that cell proliferation was already blocked at the time when LPA was added to the cells, NPCs were preincubated with HU prior to LPA addition. Preincubation with HU was effective in blocking the proliferative activity since only few EdU-incorporating cells could be detected 12 hours after exposure (Figures 2C and 2D). After 12 hours of HU incubation, LPA was added to the medium and the efficiency of the NPCs to increase their apical rosette lumen area was investigated. Non-proliferative NPCs exposed to LPA still significantly increased their apical rosette lumen area to a mean size of 131 ± 7 μm^2^ compared to control NPCs which exhibited a mean lumen size of only 57 ± 1 μm^2^. Exposure of LPA to cells that were still proliferative resulted in even larger rosette formation with an average apical rosette area of 206 ± 20 μm^2^ (Figures 2E and 2F). In conclusion, the data indicate that large rosette formation can be achieved without cell proliferation. However, since also a significant difference in apical rosette area between LPA-treated proliferative active- and inactive NPCs could be detected, it is very likely that cell proliferation contributes to increased rosette lumen sizes by providing more cells that can participate in lumen formation.

**Figure 2.**
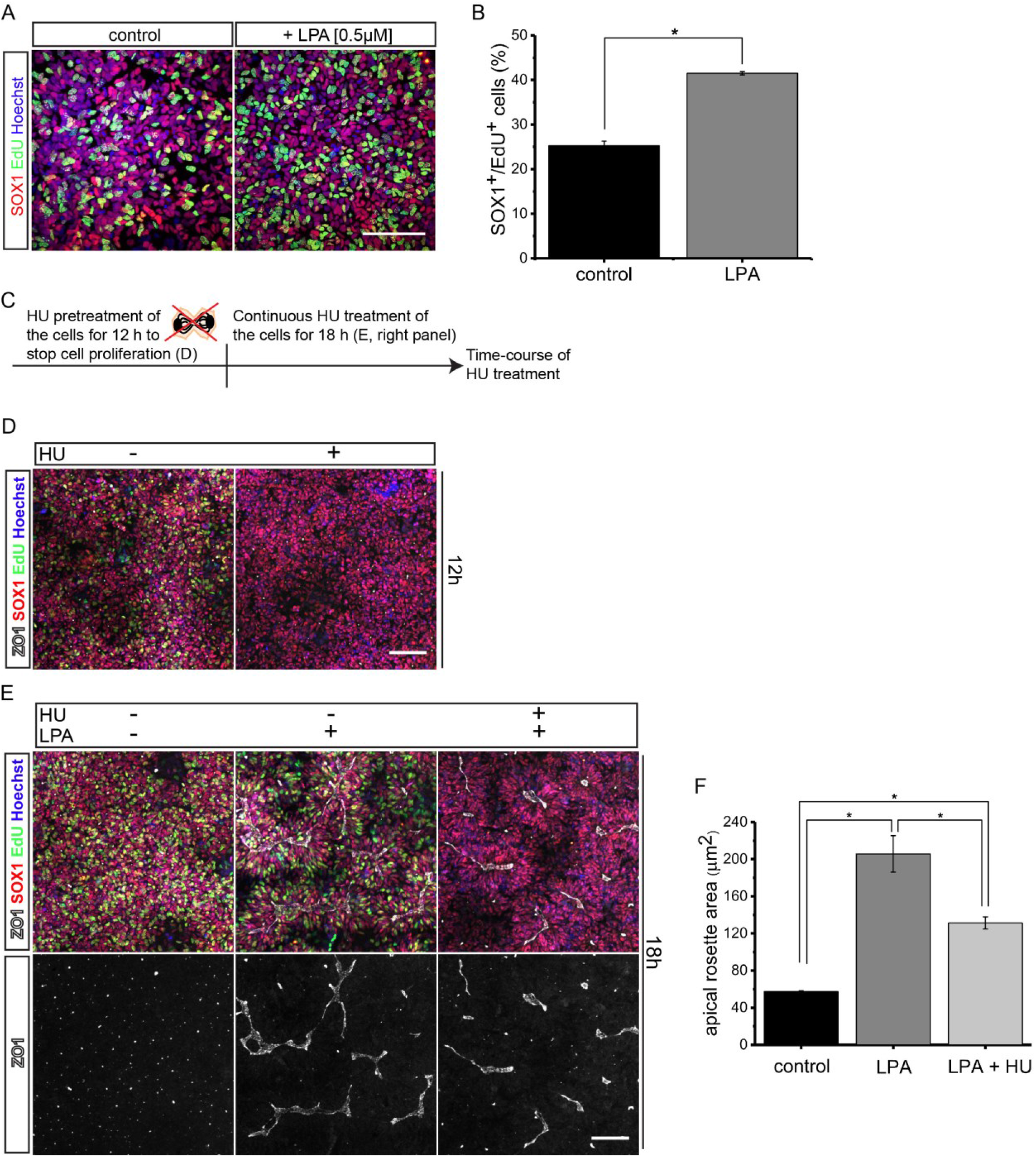
Cell proliferation is not required for large rosette formation. (A) Immunofluorescence of HESC-derived NPCs for SOX1. Prior to fixation cells were incubated for 45 min with EdU (μM). Scale bar is 100 μm. (B) Quantification of the percentage of EdU^+^/SOX1^+^ cells from the total number of SOX1^+^ cells per image. Data represent mean ± SEM. *t*-test. *= *p* < 0.001. *n* = 3. (C) Scheme depicting the experimental setup of the cell proliferation block experiment shown in D and E. (E) Immunostaining of NPCs for SOX1 and ZO-1. NPCs were cultured for 12 h in normal culturing medium (−) or in medium supplemented with 0.5 μM HU (+) and where cell proliferation was blocked. Scale bar is 100 μm. (E) Immunostaining of NPCs for SOX1 and ZO-1. NPCs were cultured in normal culturing medium (left panel), medium supplemented with 15 μM LPA (middle panel) or medium supplemented with 15 μM LPA and 0.5 μM HU (right panel). NPCs (panel A,D,E) were incubated with 30 μM EdU 45 min prior to fixation and proliferating cells were visualized using Click-iT Alexa Fluor 488. Scale bar 100 μm. (F) Quantification of mean apical rosette lumen surface area. ANOVA test followed by a Tukey a posteriori test. Data represent mean of medians ± SEM. *= *p* < 0.05. *n* = 3. Nuclei were labeled with Hoechst 33342.

### LPA induces apical domain enlargement

To better understand the changes in the epithelial organization that lead to large rosette formation of NPCs, we performed an experiment in which we initially exposed NPCs to 15 μM LPA in a volume of 0.5 ml medium and fixed the cells at consecutive time points over a period of 24 h. NPCs were fixed before (0 h), and 2 h, 10 h, 18 h and 24 h after LPA administration and immunostained for SOX1 and ZO-1 to visualize their epithelial organization. Figure 3A shows representative xy- and xz views of the NPCs at different time points after LPA addition. Prior to LPA exposure the NPCs were organized in small rosettes. ZO-1 could be detected at the apical cell sides which were constricted and enclosed a small lumen (Figures 3A and 3B, 0 h). 2 h after LPA administration the apical domains had slightly expanded in width compared to untreated controls (Figures 3A and 3B, 2 h). After 10 h, the NPCs had organized in a large and connected neuroepithelial-like cell layer in which the apical domains had adopted a much broader and unconstricted morphology and in which the individual cell-cell junctions could be easily discriminated by staining for ZO-1 (Figures 3A and 3B, 10 h). 18 h after exposure to LPA the apical domain areas started to shrink and appeared more constricted. At this point the entire morphology rather resembled a cone-shaped large rosette structure than a large connected and flat neuroepithelium (Figures 3A and 3B, 18 h). 24 h after LPA addition the cells had completely reverted to the constricted small rosette state (Figures 3A and 3B, 24 h). These observations indicate that LPA dynamically regulates the epithelial organization of NPCs within 24 h. The fact that the neuroepithelial-like morphology of the NPCs, which was observed 10 h after LPA administration, reverted over time to smaller rosettes, suggested that LPA may not be stably maintained in the culture media over time. To investigate if LPA activity within the medium gets reduced within 10 h to a level which is insufficient to induce a neuroepithelial-like phenotype in NPCs de novo, we performed an experiment in which we initially exposed NPCs for 10 h to an LPA concentration of 15 μM in a volume of 0.5 ml medium and then transferred this medium to a new culture of NPCs that had not been exposed to LPA before. Immunostaining of the first NPC culture that had been treated with 15 μM for 10 h revealed a strong neuroepithelial-like morphology (Figure 3C, upper panels). In contrast, the secondary NPCs that had been treated with medium from the first NPCs did not form larger rosettes in response to the media (Figure 3C, middle panels). As a positive control we supplemented medium that had been exposed for 10 h to NPCs with fresh LPA and added it to a parallel culture of NPCs. In this case the secondary media supplemented with fresh LPA was sufficient to induce a neuroepithelial-like phenotype with large apical domains (Figure 3C, lower panels). The results confirmed our hypothesis that LPA activity within the medium is reduced over a period of 10 h to a level where the media is not able to induce rosette enlargement. Altogether the data show that LPA promotes the organization of NPCs into a large neuroepithelial-like state with broad apical domains as long as its activity within the medium does not fall below a certain threshold (Figure 3B, dashed line). In this regard, the large rosette state that is observed 18 h after LPA administration does not represent the actual phenotype that is promoted by LPA but rather represents an intermediate state in the reversion process of the NPCs from the large apical domain-forming-neuroepithelial-like morphology back to the small rosette state as a consequence of decreasing LPA activity within the medium (Figures 3A–C).

**Figure 3.**
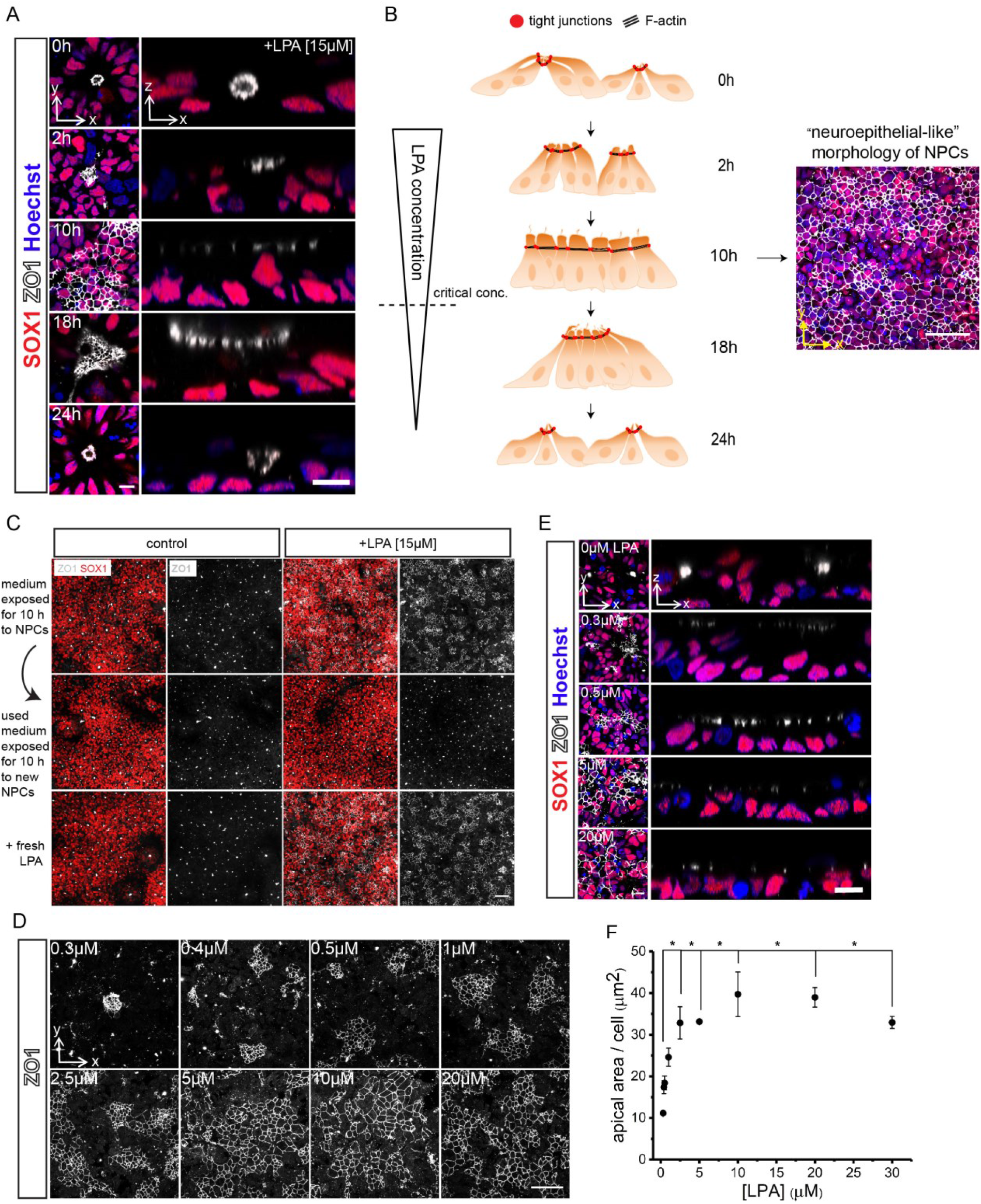
LPA dynamically regulates the epithelial organization of NPCs and increases NPC apical domain size. (A) Immunostaining of NPCs that were fixed at different time points after LPA administration for SOX1 and ZO-1. Y-x images represent top-view- and z-x side-view images. Scale bar is 10 μm. (B) Scheme depicting the changes in the epithelial organization of NPCs which were exposed to 15 μM LPA over a period of 24 h. Triangulum (left side) indicates decreasing LPA activity. Dashed line indicates the critical LPA activity inside the medium required to maintain the NPCs in a large apical domain-neuroepithelial-like state. Immunostaining shows NPCs which have adopted a large neuroepithelial-like morphology after LPA treatment. Scale bar is 50 μm. (C) Immunostaining of NPCs for SOX1 and ZO-1. Upper panels: NPCs incubated with 0.5 ml medium (upper panels, left) or medium supplemented with 15 μM LPA (upper panels, right). Middle panels: NPCs exposed for 10 h to culture medium in which NPCs had already been cultivated for 10h and that was either without-(middle panels, left) or supplemented with LPA (middle panels, right). Lower panels: Immunostaining of NPCs which were exposed to culture medium for 10 h in which NPCs had already been cultivated for 10h and that was either without-(lower panels, left) or supplemented with fresh LPA (lower panels, right). Scale bar is 50 μm. (D and E) Immunostaining of NPCs for SOX1 and ZO-1. NPCs exhibit different apical domain sizes in response to different LPA concentrations. (D) Y-x-images are top view- and z-x images are side view images. Scale bars are 100 μm (D) and 10 μm. (E) Nuclei in (A) and (E) were stained with Hoechst33342. (F) Quantification of cellular apical domain size in μm^2^. ANOVA test followed by a Tukey a posteriori test. Data represent mean of medians ± SEM. * = *p* < 0.05. *n* = 3. See also Figure S3.

### LPA increases apical domain size of NPCs in a concentration-dependent fashion

The observation that LPA promotes the arrangement of NPCs into a large neuroepithelial-like state with broad apical domains (Figures 3A and,3B), prompted us to investigate if NPC apical domain size has a dose-dependent relation to LPA concentration. This required that we expose the NPCs to stable concentrations of different LPA concentrations for at least 10 h. To achieve more stable LPA concentrations in the medium we increased the culture medium volume from 0.5 ml to 2 ml per well, thereby quadruplicating the amount of total LPA molecules per well and renewed the medium every six hours (see Experimental Procedures). Using this protocol, cells were exposed to LPA concentrations ranging from 0.3 μM to 20 μM for 36 h to allow enough time for cells to respond to the LPA. Immunofluorescence of the cells showed that the apical domains, which are regarded as the areas of low ZO-1 signal intensity in between the cell-cell borders increased with increasing LPA concentrations (Figures 3D and 3E). Quantification of the areas revealed a significant increase in the average apical domain size per cell with increasing LPA concentrations (Figure 3F). We fitted a two-state model to the experimental data (Figure S3, see Supplemental Experimental Procedures) and found that the concentration of LPA that would be necessary to achieve the 50 % of the maximal apical cell area is 0.766 μM, a concentration that is in agreement with the range of physiological LPA concentrations present in serum (Aoki et al., 2002; Yung et al., 2014). We concluded that a primary effect of LPA on the neural epithelial cells is the increase in apical domain area, resulting in the opening up of existing rosettes into an epithelial cell layer. The subsequent large rosette formation is a consequence of apical domain restriction as LPA becomes cleared from the media (see Discussion).

### Apical domain widening of NPCs in response to LPA is gene transcription dependent

Apical domain expansion- and rearrangement of epithelial cells into a large- and connected neuroepithelial-like state would be expected to be achieved by an increased lipid incorporation into the plasma membrane, growth and stabilization of the underlying cortical actin cytoskeleton and remodeling of cell junctions, such as tight junctions (Figure 4A) (Luschnig and Uv, 2014; Yeaman et al., 2004). To better understand the molecular mechanism underlying apical domain widening of NPCs we performed a microarray analysis in which we compared the gene expression of small rosette-NPCs-versus LPA-treated large rosette NPCs at 2 h, 10 h and 24h after initial exposure to LPA. Gene expression profiling of all genes that show at least an upregulation of more than two-fold in one of the three time points revealed that most upregulation occurs at 10 h, followed by 2 h and reverts to normal gene expression levels at 24 h after initial LPA addition (Figure 4B). We were wondering if the strong induction of gene expression which could be observed in the first 10 h of LPA treatment was crucial for apical domain widening of the NPCs. To investigate this we incubated NPCs for 8 h with LPA and Actinomycin D, a small molecule which is commonly used to block RNA synthesis in various cell types. Unlike NPCs which had been exposed to LPA alone and which adopted a large apical domain morphology, NPCs treated additionally with Actinomycin D failed to widen their apical domains and maintained a small rosette morphology that was identical to non-LPA treated control NPCs and control NPCs treated additionally with Actinomycin. Interestingly SOX1 expression was lost in control-Actinomycin D and LPA-Actinomycin D-treated NPCs as well which is probably a consequence of the transcriptional repression (Figure 4C). The results demonstrate that large apical domain formation of NPCs depends on an LPA-driven upregulation of gene transcription.

**Figure 4.**
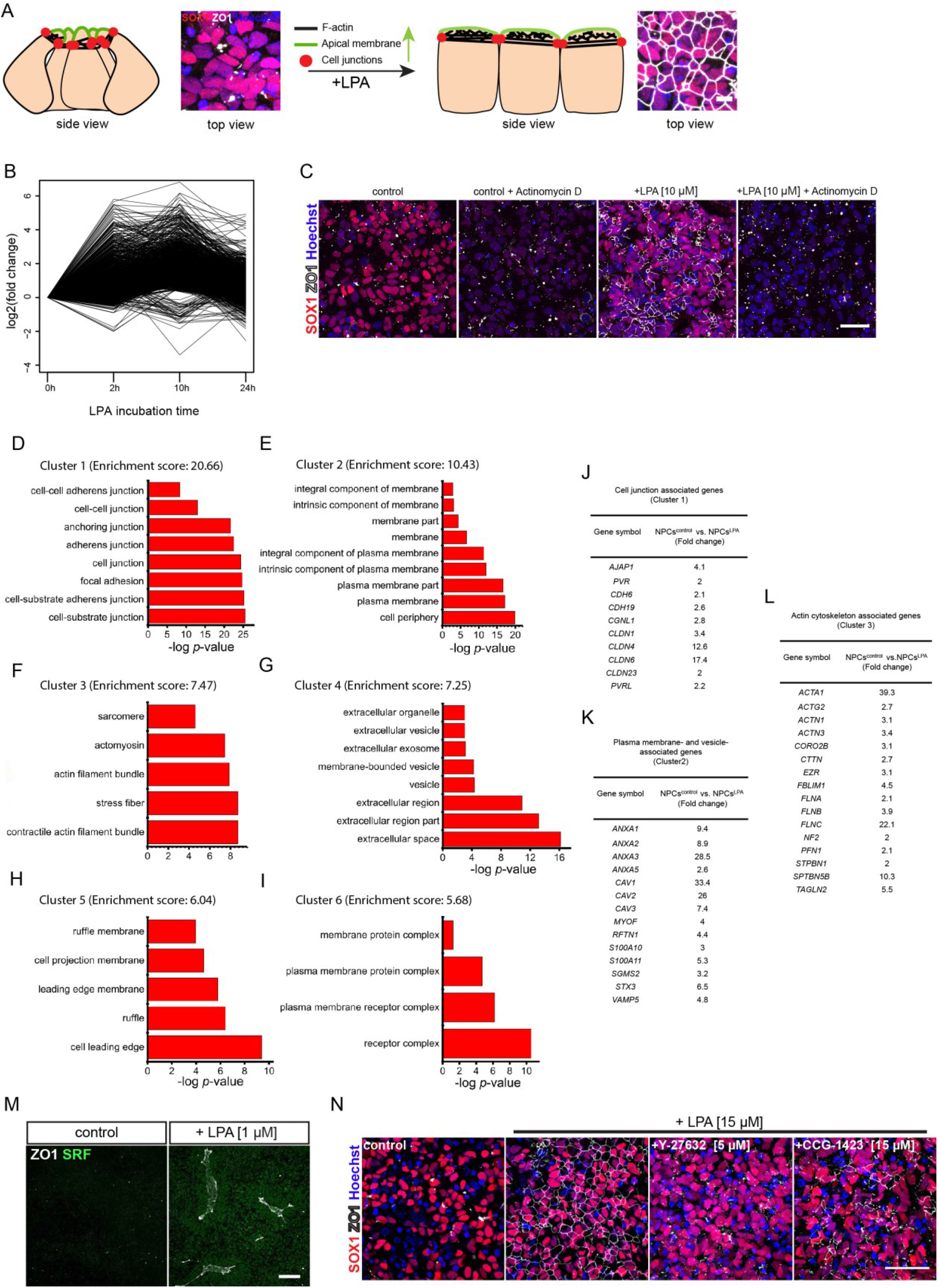
LPA upregulates genes involved in cell junction-, actin cytoskeleton and plasma membrane organization. (A) Scheme illustrating potential important cellular processes involved in apical domain increase of epithelial cells. Immunostaining of NPCs depicts the small rosettes state (left) and the large apical domain-neuroepithelial-like morphology of NPCs (+LPA, right). Scale bar is 10 μm. (B) Microarray analysis of control NPCs versus NPCs cultured in the presence of LPA at 2h, 10h and 24h after initial LPA administration. Graph depicts expression levels of all the genes that show an upregulation of more than two-fold in at least one of the three time points (log2 scale). (C) Immunostaining of NPCs for ZO-1 and SOX1 which were cultured without LPA, with LPA alone or in the presence of Actinomycin D [20ng/ml]. Nuclei were stained with Hoechst 33342. Scale bar is 25 μm. (D - I) Functional annotation clustering of all genes upregulated more than two-fold at 10 h after LPA administration. The six top GO clusters exhibiting the lowest p-value and highest enrichment of upregulated genes from the microarray are depicted. Clustering was performed using DAVID software. (J - L) Tables showing strongly upregulated genes (> two-fold) in response to LPA which are associated with cell junction-(J), plasma membrane-(K), and actin cytoskeleton organization (L). (M) Immunostaining of NPCs for ZO-1 and SRF showing the upregulation of SRF in response to LPA. Scale bar is 100 μm. (N) Immunostaining of NPCs for SOX1 and ZO-1 showing the involvement of ROCK1 and SRF in the adoption of a large neuroepithelial-like morphology. Y-276325 = ROCK1 inhibitor, CCG-1423 = inhibitor blocking SRF-associated gene transcription. Scale bar is 50 μm. Nuclei were stained with Hoechst 33342. See also Figure S4.

### LPA upregulates genes involved in apical membrane biogenesis, cell cortex and tight junction function

We used DAVID Bioinformatics software to identify cellular component gene ontology terms (GO_CC terms) that were enriched with genes that were up- or downregulated more than two-fold in the microarray data respectively. We then performed a functional annotation clustering of the GO terms to cluster GO terms that had a similar biological meaning due to sharing similar gene members. The clusters were ordered hierarchically according to their overall p-value. Clustering of the GO terms resulted in top GO clusters that were highly enriched with genes related to cell junction formation (Cluster 1), plasma membrane (Cluster 2), actin filament assembly (Cluster 3), membrane-vesicle interactions (Cluster 4), membrane movement and cell migration (Cluster 5) and plasma membrane-receptor interactions (Cluster 6) (Figures 4D–I). Among the upregulated genes that were enriched within the six top GO clusters and exhibited a differential expression of more than two-fold we found many genes related to the functions required to generate apical membrane including actin-filament assembly, cell junction-, plasma membrane- and vesicle associated genes. RNA expression levels of genes encoding for tight junction associated proteins of the claudin family (CLDN1, 4, 6, 23) and CGNL-1, the human orthologue of cingulin-like 1, were strongly upregulated (Citi et al., 1988; Furuse, 2010) (Figure 4J). Furthermore RNA expression levels of genes that are translated into caveolins 1-3(CAV1-3), which are involved in trafficking cholesterol-rich membranes that are enriched in the apical and basolateral surface were highly induced, as well as the expression of other genes involved in apical vesicle trafficking and genes related to vesicle- and membrane fusion events, such as syntaxin 3 (*STX3*) and annexin A2 (*ANXA2*) (Martin-Belmonte et al., 2007; Murata et al., 1995; Scheiffele et al., 1998; Sharma et al., 2006) (Figure 4K). Upregulation of CAV1 and CAV2 at the protein level in LPA-treated NPCs was confirmed by immunostaining and a subsequent quantification of the average amount of vesicles per cell (Figure S4). The apical surface is also characterized by a strong accumulation of cortical actin and correspondingly, a strong upregulation of RNA expression levels of actin (*ACTA1*) and actin organizing- and stabilizing genes, like profilin 1 (*PFN1*), actinin alpha 1 (*ACTN1*), cortactin (*NF2*) and genes encoding for proteins that link the actin cytoskeleton to the plasma membrane, such as ezrin (EZR), coronin2B (CORO2B) and spectrin beta 5 (SPTBN5) could be detected (Liem, 2016; Morales et al., 2004; Oikonomou et al., 2011; Rybakin and Clemen, 2005; Sohn and Goldschmidt-Clermont, 1994; Weaver et al., 2001) (Figure 4L). In contrast, GO analysis and clustering of genes that were downregulated by LPA more than two-fold mainly resulted in GO clusters related to projections, axons, synapses and calcium channels of neurons, indicating a bias in cellular components affected by LPA between the up- and downregulated genes (Table S1). The data show that LPA upregulates the expression of many genes involved in cell junction formation, actin- and plasma membrane organization which could promote the expansion of the apical domain. A full table with all the genes from the microarray enriched in the respective six top annotation GO clusters can be found in Table S2.

### Rho-/SRF signaling is involved in large rosette induction

LPA has been shown to exert many of its various cytoskeletal effects by activation of the Rho-signaling pathway and by activating the downstream transcription factor serum response factor (SRF) which is a major regulator of actin cytoskeleton dynamics (Hurst et al., 2008; Miano et al., 2007). We wanted to investigate if Rho-signaling and activation of SRF control apical domain expansion and the adoption of a larger neuroepithelial-like morphology in our NPCs. We performed immunostaining to determine if SRF is upregulated by LPA. Whereas control cells showed low SRF expression, we observed a strong signal for SRF in LPA-treated NPCs (Figure 4M). We next tested CCG-1423, a small molecule inhibitor of SRF-driven gene expression and Y-27632, an inhibitor which blocks the Rho downstream signaling factor ROCK1 on our NPCs to determine if formation of a neuroepithelial-like state in the presence of LPA could still be achieved (Evelyn et al., 2007; Watanabe et al., 2007). Immunostaining for SOX1 and ZO-1 revealed that both, application of CCG-1423 and Y-27632, resulted in a strongly disrupted ZO-1 expression pattern compared to control NPCs in which ZO-1 expression was evenly distributed around every single apical domain. The data indicate that both inhibitors could to a large extent prevent adoption of a neuroepithelial-like morphology of NPCs in the presence of LPA (Figure 4N).

### Maintenance of apical domain forming capacity in NPCs for over 52 passages

Under standard cell culture conditions, NPCs cultured in N2B27 + FGF2/EGF can be maintained for many passages without losing SOX1 expression but they maintain a small rosette organization (Figure S1J). We wanted to investigate whether NPCs maintained their inducibility by LPA at different passage numbers. Under normal culture conditions, NPCs at RP1 or RP2 formed very large rosettes in response to LPA. However at RP3 the ability of the cells to adopt a larger rosette morphology decreased and at RP4 the cells did not respond anymore to a one day pulse-treatment of LPA and remained organized in small rosettes (Figure 5A). In a next experiment, we tested if a continuous exposure of NPCs to LPA could maintain the competence to form large apical domains. We cultured NPCs from RP1 with a continuous exposure to 1 μM of LPA followed by 2 μM LPA (see Experimental Procedures) for more than 52 passages and examined the epithelial organization of the cells by immunostaining for SOX2 and ZO-1. Continuous exposure of NPCs to this level of LPA yielded SOX2^+^ cultures with strongly expanded cellular apical domains and a large rosette morphology. The maintenance of apical domain expansion was preserved for over 52 passages (Figure 5B, Figure S5A). The data suggest that LPA can maintain SOX2^+^ NPCs in a state with a strong potential to epithelialize and to form large rosettes. LPA has been reported to prevent neuronal differentiation and neurite outgrowth in iPS-cell derived neurospheres (Frisca et al., 2013). We next tested if the maintenance of a large apical domain NPC phenotype correlated with a reduced neuronal- and glial differentiation potential of the cells. We exposed NPCs directly after passaging for three days to 0.5 μM LPA. To ensure that the LPA concentration within the medium stayed constant over the entire culturing period, we increased the culturing volume to 2 ml and exchanged the medium to fresh medium every 12 h. After three days the cells were immunostained for SOX1, ZO-1 and the neuronal markers TUBB3 and HuC/HuD. Compared to control NPCs that had a small rosette morphology, the cells supplied with LPA exhibited broad apical domains and a strong neuroepithelial-like morphology with an even ZO-1 distribution all around their apicolateral regions (Figure 5C, upper panels). Immunostaining for HuC/HuD and quantification revealed that 3.9 ± 0.5 % of the SOX1^+^ cells from the control co-expressed HuC/HuD, whereas 0.9 ± 0.2 % of the LPA-treated NPCs showed co-expression (Figure 5C, middle panels, Figure 5D). Also staining for TUBB3 indicated a strong neuronal differentiation and neurite outgrowth rate in untreated NPCs compared to LPA-treated NPCs (Figure S5B). LPA has been reported to increase apoptosis in HESC-derived neurospheres (Frisca et al., 2013). To exclude the possibility that the observed reduced neuronal differentiation rates were a consequence of a LPA-stimulated increase in apoptosis in the developing neurons we stained the cells for the apoptotic marker CASP3 and compared the percentage of HuC/HuD^+^ and CASP3^+^ double positive cells in LPA-treated cultures to control NPCs. Quantification of the HuC/HuD^+^ cells expressing CASP3 did not reveal a significant difference between control NPCs and LPA-treated NPCs (17 ± 1 % in control NPCs versus 21 ± 2 % in LPA-treated NPCs), indicating that LPA does not significantly increase the apoptotic rate of early born neurons (Figures 5C, middle panels and Figure 5E). We next assessed if long-term LPA exposure could also prevent the transformation of NPCs towards a radial glial phenotype. In former publications it was shown that LPA could not significantly decrease glial differentiation in HESC- and iPSC-derived plated neurospheres (Dottori et al., 2008; Frisca et al., 2013). We compared glial differentiation of NPCs which had been cultured for 22 passages either in normal medium or medium which was continuously supplemented with 1 μM and 2 μM LPA and stained the cells for GFAP, a marker of radial glia identity. Immunostaining revealed a high GFAP signal in the control NPCs whereas the cells which had been exposed to LPA exhibited a strongly reduced GFAP signal intensity. We quantified the GFAP signal by measuring the overall GFAP intensity and normalized it to the total number of cells per image. Quantification resulted in a 92 % decrease of the GFAP signal in LPA-treated NPCs (Figures 5C, lower panels and Figure 5F). Taken together our data show that LPA maintains NPCs in a large rosette SOX2^+^ NPC state and prevents neuronal and glial differentiation.

**Figure 5.**
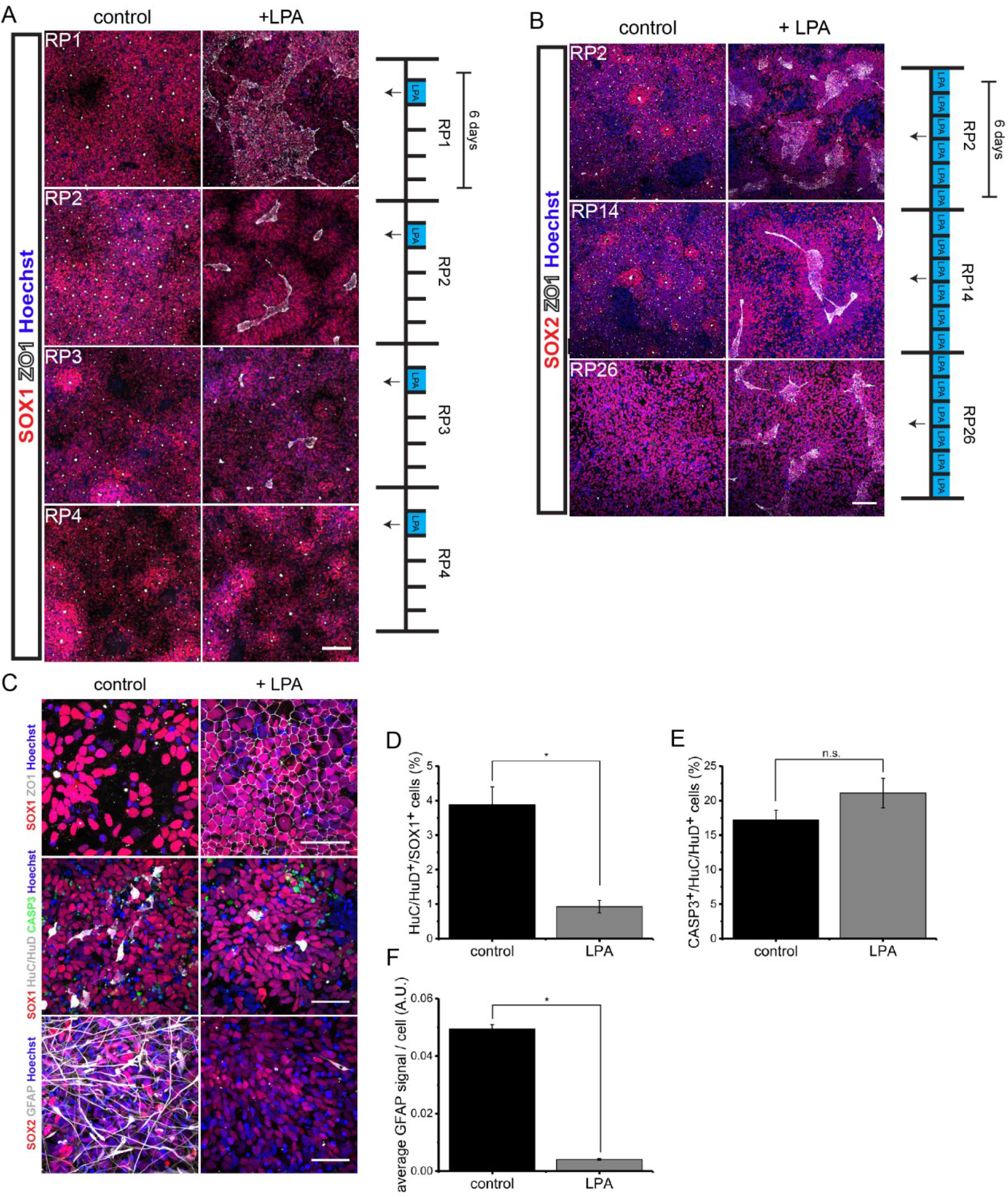
Constant LPA exposure maintains the large apical domain forming- and large rosette formation capacity of NPCs. (A) Immunostaining of untreated- or LPA-treated NPCs at different passaging numbers for SOX1 and ZO-1. Scale bar is 100 μm. (B) Immunofluorescence of NPCs for SOX1 and ZO-1. NPCs have constantly been cultured without or with LPA for 26 passages. Scale bar is 100 μm. (C) Immunostaining of NPCs for SOX1, SOX2, ZO-1, HuC/HuD, GFAP and CASP3. Scale bars are 50 μm. (D) Quantification of the percentage of HuC^+^/HuD^+^/SOX1^+^ cells from the total number of SOX1^+^ cells. (E) Quantification of HuC^+^/HuD^+^/Caspase3^+^ double-positive cells. (F) Quantification of the average GFAP intensity (AU) per cell. Nuclei were stained with Hoechst 33342. Statistical significance was analyzed using a *t*-test. All experiments were performed in triplicates. * *p* < 0.05. See also Figure S5.

## Discussion

Our results demonstrate that LPA is a conserved factor which increases neural rosette lumen size. The size increase is predominantly achieved by an increase in the individual apical domain sizes of the rosette-forming axolotl- and human NPCs but also involves an increase in cell number per rosette due to cell proliferation and merging of cells from adjacent rosettes. Large rosette formation is a very dynamic process and depends on the level of LPA activity the cells are exposed to at a given time point. At high activity LPA disrupts the small rosette organization, widens the apical domains to a maximal extent and promotes the arrangement of the NPCs into a large neuroepithelial-like cell layer in which all the cells are connected via cell junctions. In this regard the large neuroepithelial-like phenotype of NPCs which is observed 10 h after LPA administration can be regarded as one gigantic rosette with a huge lumen surface area that consists of all the apical domains of the NPCs that constitute the rosette. The gigantic neuroepithelial-like rosette morphology of the NPCs is the phenotype which is caused by high LPA activity and is maintained as long as there is enough LPA inside the medium. Our data show that LPA activity gets eliminated from the medium over time if LPA is not constantly renewed. As a consequence of reduced activity the apical domains of the cells start to shrink, the lumen size gets reduced and the gigantic rosette first collapses into several larger rosettes which gradually reduce in size and finally fully collapse back into many small rosettes. In this regard the very striking large rosette morphology which is observed 18 h after a single pulse of LPA administration does not represent the actual phenotype which is caused by high LPA activity, but rather reflects an intermediate epithelial organization of the NPCs in their reversion process back to the small rosette state due to a decreasing LPA activity in the medium.

Gene expression profiling of small rosette-forming NPCs versus LPA-treated large rosette-forming NPCs revealed a strong upregulation of gene expression in the first 10 h after LPA administration. The upregulation of transcription was essential for apical domain widening since blockage of gene transcription by Actinomycin D could prevent apical domain widening in the presence of LPA. A subsequent functional annotation clustering of the GO terms in which genes that were upregulated in response to LPA were significantly enriched revealed top GO term clusters related to cell junction organization, actin filament assembly, plasma membrane organization, vesicle-membrane interactions, membrane movement and cell migration and plasma membrane-receptor interactions. The GO terms have been described in literature as important cellular processes during luminogenesis (Datta et al., 2011; Luschnig and Uv, 2014). Strongly upregulated genes (> two-fold) found within those clusters were genes of the tight junction-forming claudin family (*CLDN1, 4, 6, 23*) and genes involved in actin filament assembly and actin filament-membrane interactions like *ACTA1, PFN1, ACTN1, SPTBN1, SPTBN5B, FLMNC* and *EZR*. Furthermore we observed an upregulation of RNA expression levels of genes involved in mediating vesicle membrane interactions and vesicle fusion. In particular the set that was identified, *STX3, ANXA2, S100A10, CAV1* and *CAV2*, is associated with biogenesis of cholesterol-rich lipid domains that are enriched in the apical domain of epithelial cells. Clustering of the genes which were downregulated by LPA revealed top GO clusters mainly related to neuron development and maturation, such as formation of neuronal projections, dendrite- and axon formation as well as synapse and potassium channel assembly. The fact that GO terms which are related to cellular processes that are important for lumen formation could only be identified in the upregulated genes suggests that those genes mainly contribute to apical domain increase and the large rosette phenotype.

From the literature it is known that deregulation of components of the Hippo signaling pathway, such as knockdown of the scaffolding protein KIBRA or expression of the CDC42 activator DBL3 can lead to an expansion of the apical domain in MDCK cells. Also overexpression of the apically localized scaffolding protein Crumbs resulted in an apical expansion of Drosophila early embryonic epithelia (Wodarz et al., 1995; Yoshihama et al., 2011; Zihni et al., 2014). Whether LPA, which is known to be an extracellular stimulator of the Hippo signaling pathway, mediates apical domain expansion by promoting the expression or downregulation of components of the Hippo signaling pathway is not yet clear. Analysis of the expression values of Hippo signaling pathway components that are known from the literature in our microarray data revealed an inconsistent picture with many stimulators of apical basal polarity establishment being slightly upregulated but at the same time others being downregulated in response to LPA. A list of known Hippo signaling components whose expression was analyzed in our microarray data can be found in (Yu and Guan, 2013). In addition to this, KIBRA expression was upregulated 1.9 fold and Crumbs levels downregulated by 3.5 fold in our microarray data. This contradicts the previous studies which have shown that knockdown of KIBRA and overexpression of Crumbs leads to apical domain expansion, indicating that LPA might act differently on the cells to promote apical domain expansion.

It is known that the binding of LPA to most of its receptors activates the Rho signaling pathway and the associated downstream factors ROCK1 and SRF. Both factors are key regulators of actin dynamics and actin-based cytoskeletal rearrangements (Miano et al., 2007; Riento and Ridley, 2003). In our experiments we show that inhibition of ROCK1 and blockage of SRF transcriptional activity could to a large extent prevent the organization of NPCs into a large neuroepithelial-like rosette phenotype in response to LPA, indicating a strong impact of actin filament-based cytoskeletal changes in large rosette formation and apical domain increase. Aliterature-based metacore analysis of all SRF target genes revealed a total of 439 binding targets. 105 of those target genes were found to be up- or downregulated (> two-fold) in our microarray data and most of the upregulated genes were found to be transcriptional activators (Table S3, Figure S6). The fact that LPA upregulated SRF in our experiments and the fact that around 25% of all SRF target genes exhibited differential gene expression in response to LPA suggests a predominant role for SRF in mediating apical domain increase and large rosette formation.

NPCs which organize in rosettes are considered to represent a neural stem cell type that can differentiate into diverse neuronal and glial subtypes in response to instructive patterning cues and thus nowadays finds application in many neuronal differentiation protocols (Broccoli et al., 2014; Kriks et al., 2011). Unfortunately the rosette state cannot be maintained for many passages as the NPCs tend to differentiate into later stage NPCs with reduced rosette formation capacity. In addition, NPCs spontaneously differentiate into neuronal subtypes which contaminate the culture. Elkabetz et al. have reported that addition of SHH- and the Notch signaling agonists Dll4 and Jagged-1 could maintain a large rosette state for at least four passages whereas NPCs that were cultured in the presence of FGF2 and EGF, two growth factors which have been shown to promote proliferation of NSCs, quickly lost rosette formation capacity. In addition Koch et al., have developed a protocol where NPCs could be passaged for > 150 passages in the presence of FGF2 and EGF without losing their small rosette organization (Elkabetz et al., 2008; Koch et al., 2009). In our experiments we show that continuous exposure of LPA to NPCs in addition to FGF2 and EGF resulted in the propagation of a very large apical domain forming neuroepithelial-like morphology of the NPCs which could be maintained for more than 52 passages. The lumen of the rosettes was much larger compared to NPCs which were cultured in FGF2 and EGF alone and which resulted in an almost complete loss of small rosette organization over time. In future, neuronal differentiation experiments need to be performed to investigate whether the maintenance of the epithelial organization in LPA-exposed NPCs at high passaging numbers has positive impact on the neuronal differentiation potential compared to control NPCs.

A speculative biological in vivo function of LPA could be to regulate the process of lumen formation and the degree of neurogenesis during neural tube development. LPA has been shown to be present inside the amniotic fluid and cerebrospinal fluid (Yung et al., 2014). In the developing mouse cortex knockdown of LPAR1 resulted in a shift of the mitotic cleavage plane from vertical towards horizontal positions in progenitor cells of the ventricular zone. Vertical cleavages generally result in symmetric cell divisions due to the equal inheritance of apical and basal membrane constituents. In contrast, during horizontal cleavages the membrane domains are inherited asymmetrically. One daughter cell inherits the basal membrane constituents and rather differentiates into a neuron, whereas the other daughter cell receives all the apical membrane constituents and stays as a stem cell (Estivill-Torrus et al., 2008; Gotz and Huttner, 2005). Hence, by regulating the degree of apical domain size, different LPA concentrations could actively be increasing or decreasing the likelihood of a symmetric or asymmetric cell division and thereby actively regulate the degree of lumen size and neurogenesis. Further experiments need to be performed to confirm whether the LPA-mediated increase in apical domains size creates a shift in the likelihood of symmetric versus asymmetric cell divisions in culture or whether the differentiation-suppressive activity of LPA acts independently of increase in apical domain size.

## Experimental Procedures

### Axolotl neural stem cell culture

Axolotl spinal cord explants were dissociated into single cells using PBS/EDTA and plated on 12-well plates coated with gelatin at a density of 32000 cells / cm^2^. Cells were passaged every 3 weeks and cultured in DMEM/F-12 medium supplemented with mouse FGF2 and B27 (Gibco). For large rosette induction the culture medium was supplemented for 3 days with either 30 % FBS or 50 μM LPA one day after passaging. After three days of serum or LPA treatment cells were cultured for another 25 days in normal culturing medium to allow for large rosette formation.

### Human ES cell culture and neural differentiation

For hESC culture, the human ES cell lines H9 and H1 were used (WiCell Research Institute). HES cells were cultured as described previously (Zhu et al., 2013). Neural differentiation was performed according to a modified version of published protocols (Khattak et al., 2015). HESC clumps were cultured for 5 days in low attachment well plates in the presence of mTeSR1 medium (Stem Cell Technologies), supplemented with 2 μM dorsomorphin and 10 μM SB431542. Under floating conditions the cell clumps formed embryoid bodies (EBs). After 5 days the EBs were transferred for 7 days to well plates coated with growth factor reduced matrigel (GFRM, Corning) and cultured in the presence of the same medium. Cells started to organize into neural rosettes during that time. Rosettes were dissociated with DPBS (w/o MgCl_2_, CaCl_2_,) / 0.1 mM EDTA) into single cells and plated at a density of 450000 cells/cm^2^ on well plates coated with GFRM. Medium was changed to N2B27 medium (Pollard et al., 2006) supplemented with human FGF2 (20ng/ml, MPI-CBG Dresden, protein facility) and mouse EGF (10ng/ml). NPCs were passaged every 6 days into new well plates (RP = rosette passaging number, indicates how often the cells have been passaged). For Figure 1A (lower panels) a different neural differentiation protocol was used. NPCs were obtained using a three-dimensional cyst-based neural differentiation protocol published previously (Zhu et al., 2013).

### LPA assays

To investigate the effect of LPA on NPCs, cells at different rosette passaging numbers were treated with different concentrations of LPA or serum. Depending on the experimental setup LPA concentrations ranging from 0.1 μM to 22.5 μM were added to a volume of 0.5 ml or 2 ml N2B27 + FGF2/EGF medium in plastic- or glass bottom 24-well plates (Corning, Greiner). NPCs were exposed for different time periods to LPA ranging from 2 h to 36 h. Long-term exposure of the NPCs to LPA (Figures 3D and 3E, Figures 5B and 5C) was performed by incubating the cells with different LPA concentrations in a volume of 2 ml medium. For the experiment shown in Figures 3D and 3E, cells were exposed to LPA concentrations ranging from 0.3 μM to 20 μM for 36 h. For the experiment presented in Figures 5B and 5C (lower panels) the NPCs were constantly cultured in the presence of 1 μM LPA for the first three days and in presence of 2 μM LPA for the last three days before passaging. For measurement of neural differentiation (Figure 5C, middle panels, Figure S5B) the cells were incubated with 0.5 μM LPA for three days. Medium change was either performed every 6 h (Figures 3D and 3E), every 24 h (Figures 5B, and 5C, lower panels), or every 12 h (Figure 5C, upper and middle panels).

### Immunostaining and microscopy

NPCs were fixed in plastic- or glass bottom 24-well plates and immunostained as described previously (Zhu et al., 2013). Cells were imaged using the confocal microscopes Zeiss LSM 700 and LSM710 (10x, 20x and 63x objective) and the Leica DMI 4000 B (63x objective). Confocal images are either single stack images or z-stack images where maximum intensity projection was performed. Phase contrast images were obtained using the Leica DM IL. Images were processed using ImageJ-, Photoshop-, and Illustrator software. A full list with all primary- and secondary antibodies is provided in the Supplemental Experimental Procedures.

### Statistics and image analysis

Image analysis was performed by using Cell Profiler (Kamentsky et al., 2011). For each experimental condition, three replicates were analyzed. From each replicate the # of rosettes per image and the distribution of apical rosette area, # of SOX1^+^ cells participating in rosette formation and the cellular apical domain size were determined. Since the data were non-normally distributed the median for each replicate was determined upon which the mean and the SEM for each experimental condition was calculated. After checking the homogeneity of variances by the Brown-Forsythe test (Brown and Forsythe, 1974), statistical comparisons were performed by a t-test or an ANOVA test followed by a Tukey *a posteriori* test when comparing two conditions or more than two conditions, respectively. A more detailed description of the two-state model which was applied in Figure 3 can be found in the Supplemental Experimental Procedures.

## Acknowledgments

We thank R. Sczech from the CRTD / TU Dresden Light Microscope Facility for help with image analysis. We also thank S. Khattak from the CRTD / TU Dresden Stem cell Facility for advise with human ES cell culture and neural differentiation. Furthermore we thank K. Goehler and R. Wegner for technical assistance with axolotl NSC cultures. We are also grateful to Julia Jarrells from the MPI-CBG Microarray DNA Facility / Dresden for expert advise and assistance. This work was supported by grants from the DFG FZ-111, EXC168, and SFB655 From Cells into Tissues to EMT. O.C. is a career researcher from Consejo Nacional de Investigaciones Científicas y Técnicas (CONICET) of Argentina and was supported by a grant of Agencia Nacional de Promoci?n Científica y Tecnológica (ANPCyT) PICT-2014-3469.

## Author Contributions

J.P.M., L.M. and E.M.T. conceived the project and conceptualized the experiments. J.P.M. performed all experiments with human ES cell-derived NSCs, acquired, analyzed and interpreted the data. L.M. and K.R. performed all the experiments with axolotl NSCs, acquired, analyzed and interpreted the data. O.C., S.O. and A.dS. analyzed and interpreted the data. O.C. performed statistical analysis, developed the two-state model and revised the manuscript. J.P.M. and E.M.T. wrote the manuscript.

